# Scaling Across Environments: Temperature and nutrition independently shape the genetics of size plasticity and morphological scaling

**DOI:** 10.64898/2026.06.10.731218

**Authors:** Shampa M. Ghosh, Isabelle M. Vea, Austin S. Wilcox, W. Anthony Frankino, Alexander W. Shingleton

## Abstract

Across animals, variation in adult body size is accompanied by coordinated variation in the size of individual morphological traits. However, the same morphological trait can scale differently with body size depending on *what* drives the size variation. In *Drosophila melanogaster*, for example, wing size scales differently with body size when size varies because of developmental nutrition versus developmental temperature. Whether the genetic basis of size plasticity and scaling is shared across different environmental regulators of size remains unclear, but is central to predicting how selection acts on the developmental mechanisms that regulate trait size, plasticity and morphological scaling. Using ~200 isogenic *D. melanogaster* lineages, we measured wing and leg size across nutritional and thermal treatments. For each lineage, we estimated nutritional and thermal plasticity for both traits, as well as the wing-leg individual-level scaling relationship, or ILSR, generated by each environmental source of size variation. We found extensive genetic variation in both thermal and nutritional plasticity for wings and legs, and in the slope of the ILSR between them. However, a lineage’s thermal plasticity was genetically uncorrelated with its nutritional plasticity for either trait, and we detected no genetic correlation between the slopes of thermal and nutritional wing-leg ILSRs. We also found no genetic correlation in the slope of nutritional wing-leg ILSRs across temperatures. Thus, the slope of a lineage’s nutritional ILSR at 17°C was not predictive of its slope at 25°C of 28°C. Nevertheless, the overall pattern of nutritional ILSRs was conserved across temperatures. These results suggest that the genetic architecture of size plasticity and scaling depends on the environmental source of size variation. Consequently, the evolutionary response of scaling to selection in heterogeneous environments may not be predictable from genetic variation measured in any single environment.

## Introduction

Variation in adult body size within populations is perhaps one of the most familiar forms of phenotypic variation and reflects the combined effects of genetic and environmental factors on growth and development. To maintain function, variation in body size must be accompanied by coordinated variation in the sizes of individual-level morphological traits. Thus, larger humans tend to have longer legs, arms and torsos, bigger livers and larger hearts. The resulting covariation in trait size among adults within a population is known as a *static morphological scaling relationship* and defines the characteristic shape of that population or species (Gould 1966; Klingenberg and Zimmermann 1992; Stern and Emlen 1999; Shingleton et al. 2007; Shingleton 2010). Indeed, to a first approximation, the evolution of morphology is the evolution of scaling (Shingleton 2010). For this reason, scaling relationships have been studied intensely for more than 150 years (Gayon 2000). Nevertheless, the mechanisms governing their evolution remain poorly understood.

Morphological scaling relationships are typically represented as log-linear relationships between trait sizes, described by the allometric equation: log *y = α log x +b*, where *x* and *y* are the sizes of individual-level traits, measured on the same scale (Huxley and Tessier 1936). Here, *α* (the allometric coefficient) describes the extent to which the size of *y* changes relative to *x* as both vary with overall body size, while *b* captures the relative size of the two traits at a reference size (log *x* = 0). The equation describes observed relationships between trait sizes without assuming any underlying causes. Nevertheless, the allometric coefficient reflects the extent to which variation in the size of trait *x* is accompanied by variation in the size of trait *y*, and thus captures the relative sensitivity of the two traits to the factors that generate size variation.

Importantly, the allometric coefficient for the scaling relationship between two traits can vary depending on the factor that generates size variation. For example, in *Drosophila melanogaster*, the allometric coefficient for the scaling relationship between wing and body size is ~1 when size varies due to nutritional variation during larval development (‘developmental nutrition’), but ~1.6 when size varies due to developmental temperature during the same ontogenetic period (Shingleton et al. 2009). For the leg, in contrast, the same coefficients are ~1 and ~0.6, respectively. If trait size were determined mechanistically by body size, then the allometric coefficient would be invariant across different environmental affectors of size. That this is not the case indicates that traits differ in their sensitivities to different factors that regulate size, and that overall body size is a reflection of the aggregate of these sensitivities. Scaling relationships therefore emerge from how individual-level traits respond to growth-regulatory signals rather than representing fixed properties of the organism as a whole.

This distinction impacts how scaling relationships are interpreted. The scaling relationship observed in a natural population, referred to as a *Population-Level Scaling Relationship* (PLSR), reflects the collection of genetic and environmental factors that determine trait size in each individual-level (**Figure 1A**). Because individual-levels differ genetically and are subject to different developmental environments, a population-level relationship is a statistical summary of myriad individual-levelly-unique developmental processes rather than the expression of a single mechanism. As such, a population-level relationship cannot reveal directly the genetic variation upon which selection acts to change scaling, since selection operates on growth and developmental processes *within individuals*, not on the PLSR itself. Problematically, this means that the pattern or form of selection that produces/favors a particular scaling relationship cannot be inferred from the pattern of scaling exhibited in a population. Two statistically-indistinguishable PLSRs may arise from different patterns of growth regulation by the same developmental mechanisms or from different growth-regulatory processes entirely (Dreyer et al. 2016). Further, they may have evolved convergently in response to different modes of selection, and, correspondingly, may respond differently to the same form of selection (Dreyer et al. 2016). Explaining the evolution of scaling therefore requires an understanding of scaling at the level of individual-level developmental responses rather than population-level patterns alone.

**Figure 1.**
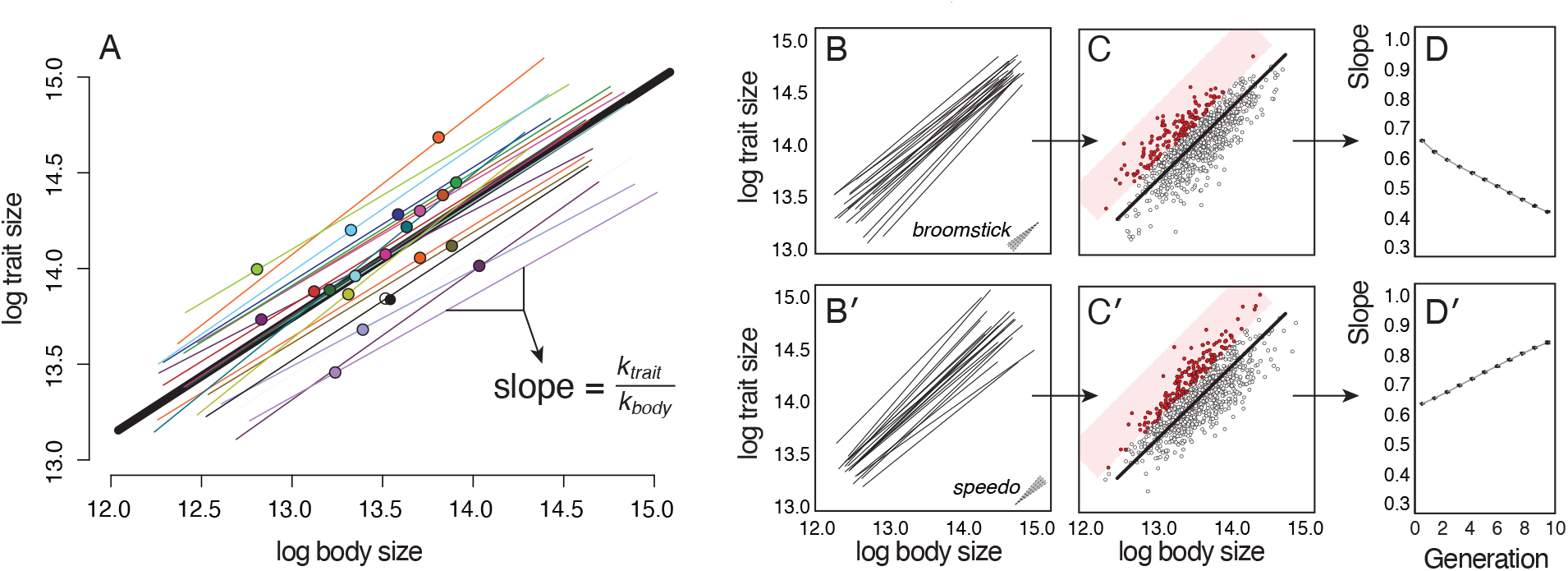
Population- and individual-level scaling relationships. (A) A population-level scaling relationship (PLSR: heavy gray line) is fit across individual-levels (colored points), each of which sits on its own individual-level scaling relationship (ILSR:colored lines). The slope of an ILSR is the ratio of the sensitivities (*k*) of the two traits to the environmental or genetic factor that generates size variation. (B, B′) The pattern of ILSRs relationships in a population determines how the PLSR responds to selection. Two populations can differ in this underlying pattern even when they share the same PLSR, here illustrated by a “broomstick” pattern, in which ILSRs fan out as they approach the origin (B), and a “speedometer” pattern, in which they fan out from the origin (B′). (C, C′) Realized individual-levels from the two populations (points) generate statistically indistinguishable PLSRs (heavy gray line), but the same pattern of selection on relative trait size (red points) targets very different subsets of ILSRs in each. (D, D′) As a consequence, the same form of selection drives the slope of the PLSRs in opposite directions across generations: the slope decreases in the broomstick population (D) and increases in the speedometer population (D′). Figure adapted from Dreyer et al. (2016).

To address this challenge, the concept of an *Individual-Level Scaling Relationship* (ILSR) has been developed (Dreyer et al. 2016; Houle et al. 2019). ILSRs describe scaling between traits for a single genotype when they covary in response to a single size-affecting factor (**Figure 1A**). Individual-level scaling relationships are obtained by rearing individuals with a single genotype across a gradient of an environmental variable (an *environmental* ILSR) (Shingleton et al. 2009), or by introducing allelic variation at a single size-regulatory genetic locus (a *genetic* ILSR) (Pavlicev et al. 2011). The slope of an ILSR (*α*) is therefore controlled by the relative sensitivity of traits to the size-influencing factor (**Figure 1A**). When the regulator is environmental, these sensitivities correspond to the environmental responsiveness of the traits, that is, their plasticity. Because the slope of an environmental ILSR captures the relative plasticity of two traits, it is this form of ILSR that we focus on here.

In practice, ILSRs can only be generated in plants or animals that can be genetically replicated, such as clonal species or experimentally produced isogenic lineages. Consequently, for most species, ILSRs are cryptic, with the observed trait sizes being single points on unseen ILSRs, each ILSR generated by the individual’s genotypic response to a single size-regulatory factor (**Figure 1A)**. Although these relationships are typically unobserved, their variation reflects genetic variation in the developmental mechanisms that co-regulate trait growth in response to a particular environmental factor. It is this variation, rather than the PLSR itself, upon which selection acts to change morphology.

Mathematical modeling of ILSRs reveals that their pattern dictates how a PLSR will respond to selection (**Figure 1B**). Because the pattern exhibited by ILSRs in morphospace reflects genetic variation in the relative sensitivities to growth-regulatory environmental factors among individual-levels, a form of GxE interaction, the pattern determine the direction and magnitude of evolutionary response of scaling to selection. Critically, the pattern of ILSRs cannot easily be inferred from the corresponding PLSR (**Figure 1C**). Thus two populations may have ostensibly identical PLSRs, but different patterns of variation in growth regulating mechanisms, and thus can respond in different ways to the same selective pressure (**Figure 1D**).

Despite their centrality to understanding the evolution of scaling, few studies have described the pattern of individual-level scaling relationships in a population. Those that have typically examine responses to a single environmental factor, most often nutrition. Although different environmental factors produce distinct ILSRs, it is not known whether the pattern of ILSRs that are generated in response to one environmental factor (for example nutritional ILSRs), changes in response to a second environmental factor (for example temperature), a form of (G x E_1_)x E_2_ interaction. This is an important question: Since the pattern of ILSRs in a population determines how it will respond to selection, environmentally-induced changes in this pattern may cause the same selective pressure to produce different evolutionary changes in scaling under different environmental conditions.

Here we determine if genotypes respond to variation in different environmental factors similarly or differently. Using multiple isogenic lineages of *Drosophila*, we quantify the effects of temperature and nutrition on wing size, leg size, and on the ILSR between them. Because ILSRs reflect relative trait size plasticity, we first ask whether the plastic responses are genetically correlated between traits within environmental gradients and within traits between environmental gradients. We then test whether the slope of the wing-leg ILSR is genetically correlated between environments, and whether this impacts the pattern of nutritional ILSRs at different temperatures.

## Materials and Methods

All data and the R code used to analyze them are provided on Dryad (DOI: 10.5061/dryad.8sf7m0d36). The phenotypic measurements of body size under starvation treatments were originally collected and described in Vea et al.(Vea et al. 2023). All other data have not been previously reported.

### Fly Stocks

All individual-levels analyzed in this study were drawn from the Drosophila Genome Reference Panel (DGRP), a collection of approximately 200 fully sequenced, inbred, and genetically isogenic Drosophila lines derived from a single natural population originally sampled in Raleigh, North Carolina, USA (Mackay et al., 2012). Flies were reared on standard cornmeal–molasses medium (Frankino et al., 2019) under a 12:12 hour light:dark photoperiod and at 75% humidity, unless otherwise stated.

### Starvation treatment

*Drosophila* egg collection, rearing, and phenotyping followed our established protocols (Stillwell et al. 2011; Stillwell et al. 2016; Frankino et al. 2019). For each DGRP lineage, females oviposited for three days. At 24h, and 48h, eggs were collected, divided into lots of 50 and placed into vials containing 10ml of standard cornmeal molasses medium, and reared at 22°C 12:12 L:D. This generated two age cohorts of flies (D0 and D1). When third instar larvae from D0 began to pupariate, larvae from all cohorts were removed from the food and placed into empty food vials with a wet cotton plug to provide moisture. Pupae were removed from these vials and transferred to individual-level 1.5ml Eppendorf tubes, each with a small hole in the lid, to complete development to adulthood. Larvae in the D0 cohort were starved for between 0-24h before pupariation, larvae in the D1 cohort were starved for between 24-48h before pupariation. Because larvae stop feeding ~24h before pupariation (Testa et al. 2013), D0 larvae were essentially allowed to feed *ad libitum* and more-or-less achieved full adult body size. In contrast, D1 larvae were starved before larval wandering, reducing adult size depending on their size at initiation of starvation. Across all cohorts, our starvation treatment therefore generated nutritionally-induced variation in body size. We refer to D0 flies as *fed* and D1 flies as *starved*. Newly eclosed adults were collected for subsequent analysis.

### Thermal Treatment

For each DGRP lineage, three replicate parental vials were established. Females in each vial were allowed to oviposit for 24 h until at least 50 eggs were deposited. Each vial was then assigned to one of three developmental temperatures (17°C, 25°C, or 28°C). Eggs and larvae were allowed to develop at their assigned temperature until pupation and adult eclosion. Newly eclosed adults were collected for subsequent analysis.

### Body and Trait Size Measurement

Sex is a major determinant of both body size and body size plasticity in Drosophila. To reduce the number of confounding variables in our analyses we therefore measured and analyzed trait size in male flies only. Traits sizes were measured using established protocols (Shingleton et al. 2009; Stillwell et al. 2011). Briefly, *Drosophila* adults were dissected, and their right wing and right first leg mounted in dimethyl hydantoin formaldehyde (DMHF). Wing area and femur length (a proxy for leg length; (Shingleton et al. 2009) were measured a via semi-automated custom software (Metamorph, Molecular Devices LLC) that captures images from a digital camera-equipped microscope (Leica DM6000B, Leica Microsystems Inc). Femur length was squared to put it in the same dimension as wing area, and all measurements were log transformed to ensure scale invariance across traits of different sizes.

### Statistical Analysis

All analyses were conducted in *R*, and the data and script to analyze them are provided on Dryad.

A major challenge to analyzing genetic variation and covariation in plasticity and scaling, is that both are characteristics of groups rather than individual-levels. One solution is to extract summary statistics from each genotype, (for example mean trait size under each environmental condition) and use these in subsequent tests of genetic correlation. However, these summary statistics do not capture the uncertainty of measurements within each lineage, which will lead to anti-conservative estimates of statistical significance when genetic correlations are calculated from indices derived on these estimates. To address this uncertainty we first used MCMCglmm to generate 1000 independent estimates of lineage-specific trait sizes under each environmental condition (fed v. starved at 22°C and fed at 17°C, 25°C, and 28). Because, the data for the effects of nutrition and temperature on trait size were collected in two separate experiments, they were analyzed separately. The specific models were:

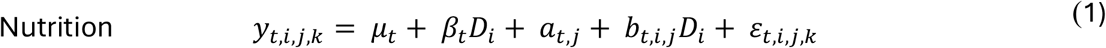

where *y*_*t,i,j*,k_ is the size of trait *t* for individual-level *k* from lineage *j* reared in condition *i, μ*_*t*_ is the population mean for trait *t* (wing or leg) in fed conditions, *β*_*t*_ is the population-level effect of starvation on trait *t, D*_*i*_ is an indicator variable (fed: *D*_*i*_ =0; starved: *D*_*i*_ =1), *a*_*t,j*_ is the random deviation of lineage *j* from the population mean size, *b*_*t,i,j*_ is the random deviation of lineage *j* from the population-level plasticity, and *ε*_*t,i,j,k*_ is random error.

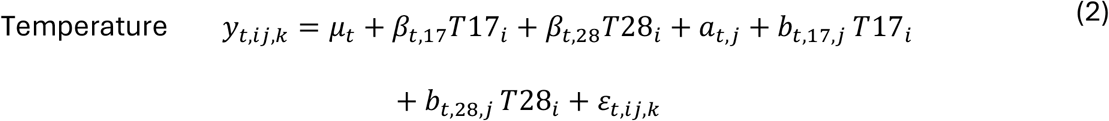

where *β*_*t*,17_ and *β*_*t*,28_ are the population-level effects of 17°C and 28°C relative to 22°C on trait *t,T*17_i_ and *T*28_i_ are indicator variables, and *b*_*t*,17,*j*_, *b*_*t*,28,*j*_ are the corresponding lineage-specific deviations. Population-level thermal plasticity between 17°C and 28°C is *β*_*t*,28_ − *β*_*t*,17_, and lineage-specific thermal plasticity is (*β*_*t*,28_ + *b*_*t*,28,*j*_) − (*β*_*t*,17_ + *b*_*t*,17,*j*_). Convergence was assessed by visual inspection of trace plots and by calculating effective sample sizes, which exceeded 200 for all parameters, indicating adequate chain mixing.

For each lineage, we extracted 1000 posterior draws of three quantities from each model: (**i**) trait size under each environmental condition (wing and leg size in fed and starved conditions from the nutrition model; wing and leg size at 17°C, 25°C, and 28°C from the temperature model); (**ii**) trait plasticity (the environmental effect size, β_t_+*b*_*t,j*_ for each trait between environmental conditions), and; (**iii**) the slope of the ILSR between wing and leg size, when they covary in response to the same environmental factor. All ILSRs were fit using major axis (MA) regression, and the slope was estimated as the ratio of the wing and leg plasticities within each environmental shift (fed – starved; 17°C – 25°C; 25°C – 28°C; 17°C – 28°C) (see Supplementary Methods for justification). ILSRs were only estimated for lineages where the 95% HPD interval of the plasticity estimates excluded zero; that is, where there was a demonstratable effect of the environment on trait size.

We then used these draws in subsequent analysis of genetic variance, covariance, and correlation within and among lineages for trait size, plasticity and scaling. For correlations of plasticity and scaling between environmental treatments, where nutritional and thermal plasticity were estimated from separate models, we computed the across-lineage correlation for each draw, pairing draws by index across models. Correlations among lineages were visualized using MA regression.

Throughout we report the model value of the summary statistics based on the 1000 posterior draws along with their 95% Highest Posterior Density (HPD) interval. For all plots, we present the 95% HPD interval of the MA, along with the modal values for each lineage. 95% HPD intervals not overlapping with zero were used to indicate strong evidence of effect.

To estimate within-lineage variance in wing and leg size at each temperature we fit the temperature model (Equation 2) using *glmmTMB*, which allows us to model temperature-specific residuals. We estimated within-lineage variance from the residual dispersion component of the fitted model, and the among-lineage variance at each temperature from the lineage variance-covariance matrix. The proportion of variance that could be attributed to within lineage variation in size at each temperature was calculated as the residual variance divided by the total variance. To test whether this variance was significant, we compared models in which the residual dispersion was either constant or allowed to vary across temperatures.

To test whether the slope of the within-temperature wing-leg ILSRs changed systematically with temperature, we used a bootstrap procedure that accounted for estimation uncertainty in the per-lineage slopes. For each of 5,000 bootstrap replicates we resampled wing and leg measurements within each lineage-temperature combination with replacement, and calculated the slope of the MA regression of wing on leg. We then fit a fixed effects model of slope in response to temperature (categorical factor), with lineage as a fixed factor, to isolate within-lineage changes across temperatures rather than variation among lineages. For each resample, we calculated the change in slope from 17°C to 25°C (δ_17.25_) and from 25°C to 28°C (δ _25.28_). Two-tailed P values for δ_17.25_ and δ _25.28_ were calculated as twice the smaller of the proportion of the 1,000 bootstrap estimates above and below zero. We used a similar bootstrap approach to determine whether there was a genetic correlation in the slope of the within-temperature wing-leg ILSR between 17°C and 25°C and between 25°C and 28°C. Because the slope of an MA regression can become unstable when the correlation between *x* and *y* is low, we only conducted analyses of within-temperature scaling on lineages where *r*_wing,leg_ > 0.3 within all three temperatures.

We also tested whether there was random (rather than systematic) variation in the effect of temperature on the slope of the wing-leg ILSR by comparing two models using a likelihood ratio test:

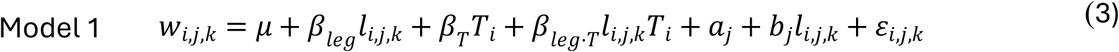

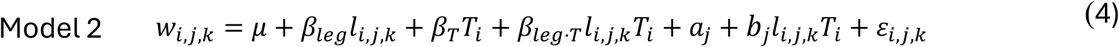

where *w*_*i,j,k*_ is wing size of individual-level *k* from lineage *j* at temperature *i, l*_*i,j,k*_ is leg size, and *T*_*i*_ is temperature. Model 1 allows random lineage-specific deviation from the average wing-leg ILSR slope (*b*_*j*_*l*_*i,j,k*_) that is constant across temperatures. Model 2 allows random lineage-specific deviation from the average wing-leg ILSR slope (*b*_*j*_*l*_*i,j,k*_*T*_*i*_) that varies with temperatures. Thus model 2 allows each lineage to respond differently to temperature with respect to the effect of temperature on wing-leg ILSR slope changes.

To determine the pattern of within-temperature ILSR we fit an MA regression to wing on leg size in each lineage at each temperature and calculated the median point of intersection (MPI) among individual-level scaling relationships ((Dreyer et al. 2016) and the bivariate mean of wing and leg size across lineages.

## Results

### There is extensive genetic variation in nutritional and thermal size plasticity of wing and leg

Both wing and leg size responded strongly to diet and temperature, but the magnitude of that response varied substantially among DGRP lineages. There was considerable genetic variation among lineages in both thermal and nutritional size plasticity (GxE) of wing and leg, with the 95% HPD interval for *V*_*GxE*_ excluding zero in all cases (**Figure 2, Supplementary Figure S1**). Consistent with previous studies (Shingleton et al. 2009), wing size was considerably more thermally plastic than leg size: the difference in thermal plasticity between the wing and the leg had a mode of −0.276 (95% HPD: −0.298, −0.256). In contrast, the leg was marginally more nutritionally plastic than the wing (differences in nutritional plasticity, mode = 0.021, 95% HPD interval: 0.007, 0.031)

**Figure 2.**
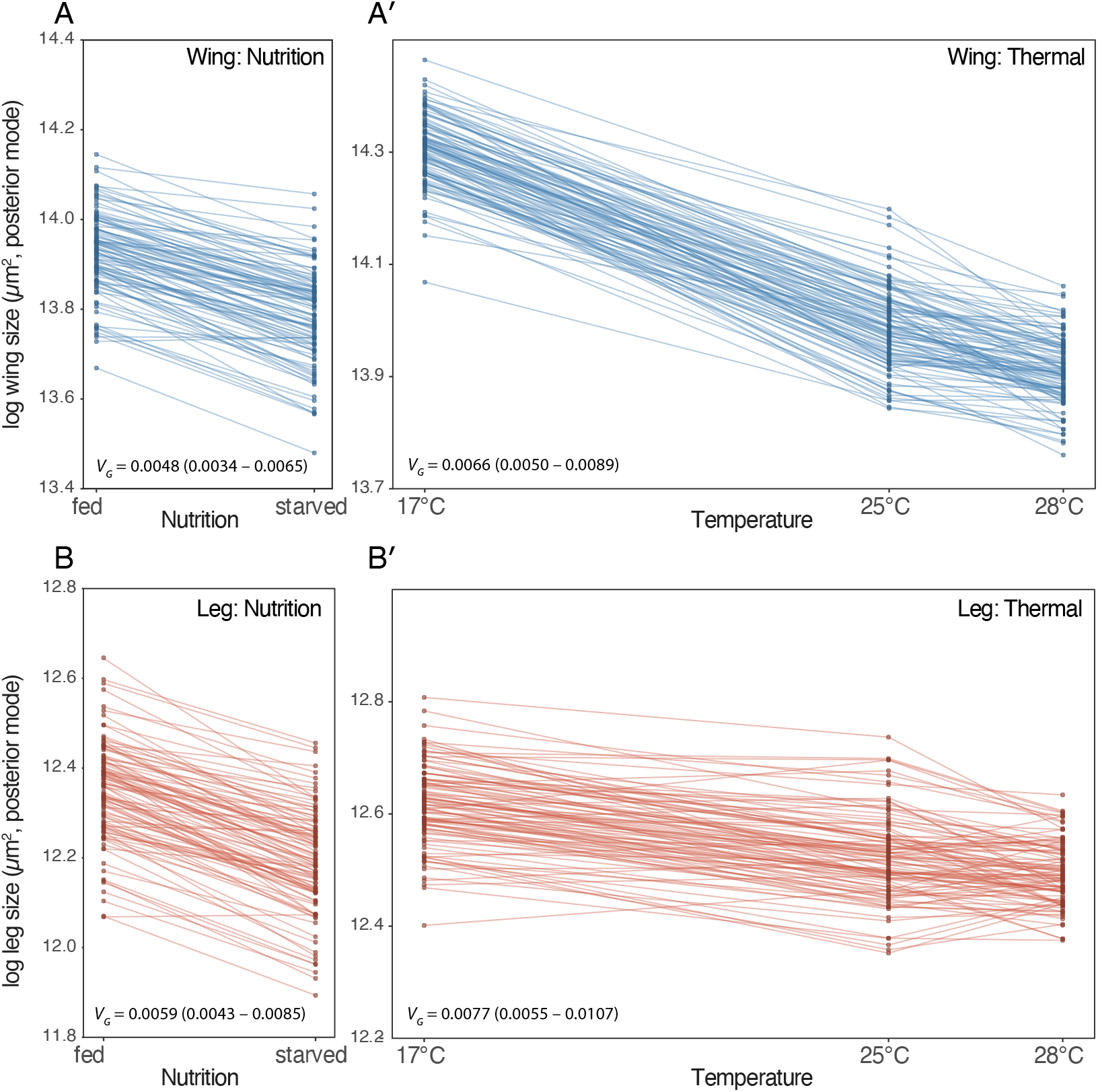
Reaction norms of wing and leg size against nutrition and temperature. Points are posterior modes for each lineage, across 1000 draws of the MCMCglm model. Note that the *y*-axes are on different scales between environments but the same scale within.

Plasticity varied among genotypes both independently of, and in proportion to, overall trait size. Previous authors have used a cross-environment genetic correlation of <1 as an indicator of genetic variation in plasticity (Via and Lande 1985; Choy et al. 2024), and for both traits across all environmental shifts (e.g. fed→starved; 17°C→25°C, etc.), this correlation was <1 (**Supplementary Table 1**). It is important to note, however, that a genetic correlation of less than 1 is a sufficient but not necessary indicator of genetic variation in plasticity of trait size. If the genetic correlation for trait size between environments =1 but the slope of that relationship differs from 1, plasticity varies among genotypes, albeit systematically with overall trait size (**Supplementary Figure S2**). For most cross-environment relationships, the slope of the relationship differed from 1 (**Supplementary Table 1**), indicating that trait plasticity either increased or decreased with overall size, although not consistently in direction.

### Thermal and nutritional plasticity are not genetically correlated

A lineage’s sensitivity to temperature was genetically independent of its sensitivity to nutrition. Our data strongly supported a genetic correlation between the plasticity of wing size and plasticity of leg size, as long as both are responding to the same environmental variable (fed – starved; 17°C – 25°C; 25°C – 28°C; 17°C – 28°C, **Supplementary Table 2**). In contrast, we found no evidence that thermal plasticity is genetically correlated with nutritional plasticity for either trait (**Figure 3**): In all cases, the 95% HPD intervals for the correlations contained zero (**Supplementary Table 2**).

**Figure 3.**
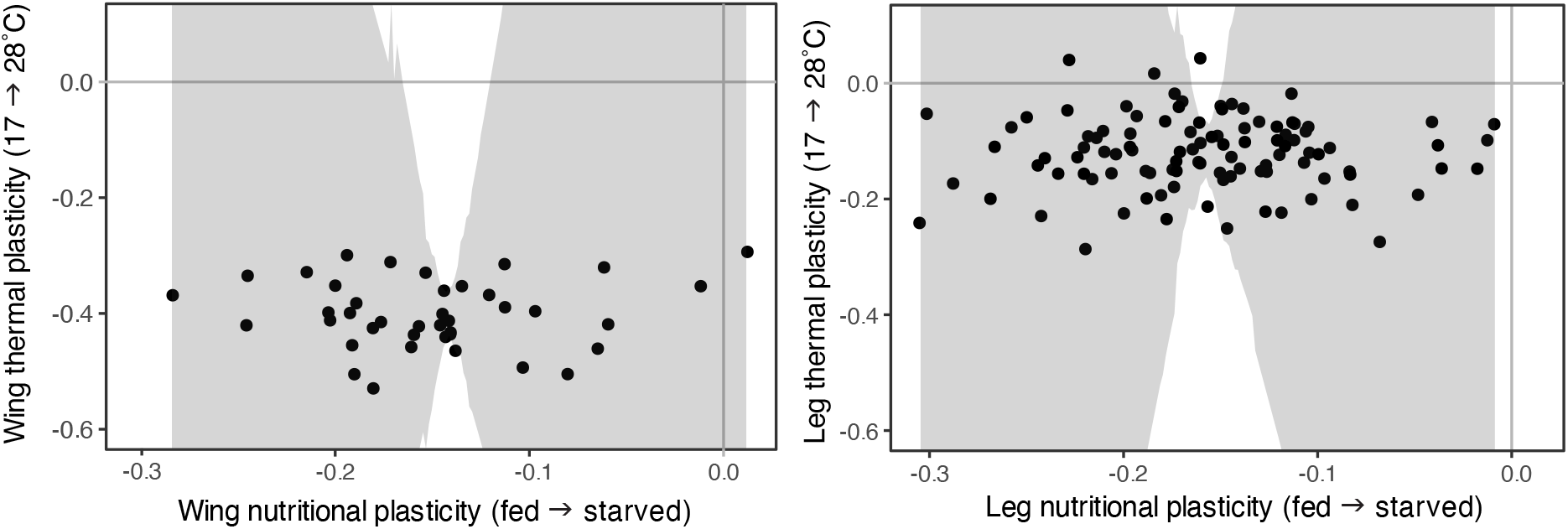
There is no genetic correlation between thermal and nutritional plasticity within traits. Points are modal values of plasticity estimates per lineage. The HPDs (grey) for the correlation between nutritional and thermal plasticity contained zero for both wing and leg (**Supplementary Table 2**).

There was considerable genetic variation in the shape of the thermal reaction norms for both the wing and the leg. For both traits, the data strongly supported a negative correlation in plasticity between the 17°C–25°C and 25°C–28°C intervals (**Supplementary Table 2**): lineages that shrank the most between 17°C and 25°C shrank the least between 25°C and 28°C, and vice versa. This is consistent with the thermal reaction norms being non-linear across the full temperature range, but where they converge at the temperature extremes (17°C and 28°C) and fan out at intermediate temperatures (25°C) (**Figure 2**).

### Thermal and nutritional scaling relationships are not genetically correlated

Although we found no evidence that size plasticity is correlated across different environmental regulators of size, it remains possible that the slope of ILSRs between traits, which reflects relative trait plasticity, is genetically correlated across environments. That is, lineages with a steeper wing-leg scaling relationship when size varies with nutrition (nutritional ILSRs) may also show a steeper wing-leg scaling relationship when size varies with temperature (thermal ILSRs). However, our data did not support this hypothesis (**Figure 4**): In all cases, the 95% HPD intervals for the correlations for the slope of wing-leg ILSRs between different environmental regulators of size contained zero (**Supplementary Table 3**).

**Figure 4.**
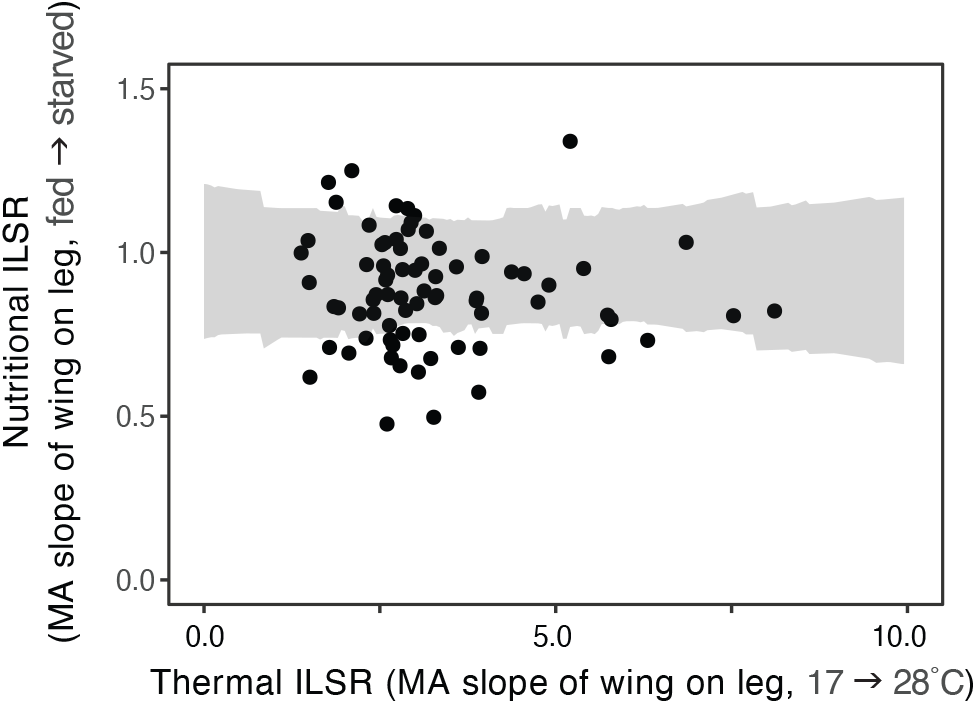
There is no genetic correlation between the thermal and nutritional ILSRs for wing against leg size. Points are modal values of scaling estimates per lineage. The HPD (grey) for the correlation between nutritional and thermal scaling contains zero (**Supplementary Table 2**).

### Nutritional plasticity changes with temperature but scaling does not

The observation that scaling relationships generated by different environmental factors are not genetically correlated is consistent with previous studies that show that the pattern of scaling is different when body size varies in response to nutrition versus temperature (Shingleton et al. 2009). These data suggest that the developmental mechanisms that regulate scaling in response to variation in temperature are independent to those that regulate scaling in response to variation in nutrition. If true, then we would expect nutritional scaling to be invariant across temperatures, and vice versa.

Testing this prediction requires measurement of the nutritional scaling relationship at different temperatures. Although we did not explicitly impose variation in nutrition at different temperatures in our experiment, 30-61% of the variation in trait size across lineages and temperatures could be attributed to within-lineage variation. This within-temperature variation was a consequence of nutritional variation, evident from the observation that a lineage’s wing-leg ILSR within a thermal treatment aligned with that lineage’s nutritional ILSR, as measure using our diet manipulation (**Supplementary Figure S3**). We therefore used the within-temperature variation in trait size as a measure of a lineages’ nutritional plasticity, and the MA regression of wing on leg size within a temperature as the nutritional ILSR for the two traits at that temperature; that is, their relative nutritional plasticity.

Temperature systematically affected the *magnitude* of nutritional plasticity across lineages. Within-lineage nutritional plasticity differed across temperatures for both traits (leg: χ^2^_2_ = 43.54, P < 0.0001; wing: χ^2^_2_ = 7.55, P = 0.023), increasing between 25 °C and 28 °C for the wing (glmmTMB, z = 2.36, P = 0.018) and between 17 °C and 25 °C for the leg (glmmTMB, z = 4.87, P < 0.0001). Thus there is a (GxE_1_)xE_2_ effect on the size of traits, such that a lineage’s response to variation in nutrition depends on temperature.

Temperature did not, however, systematically affect *relative* nutritional plasticity. Across lineages, the slope of the wing–leg nutritional ILSR showed no consistent directional change with temperature (two-tailed bootstrap P-value, δ_17.25_ = −1.43, *P* = 0.61; δ_25.28_ = 1.40, *P* = 0.50). Temperature therefore appears to affect the overall magnitude of nutritional plasticity without consistently changing how plastic the wing is relative to the leg.

This does not mean slope was unaffected by temperature, but rather that its response was lineage-specific rather than shared. A model of wing size allowing the temperature effect on slope to vary among lineages fit the data significantly better than one in which temperature acted uniformly (R^2^: 0.957 vs 0.941, respectively; likelihood-ratio test: χ^2^ = 1201.9, df = 7, P < 2.2 × 10^−16^). Consistent with this, slopes were uncorrelated across lineages between temperatures (17 °C vs 25 °C: r = 0.009, P = 0.94; 25 °C vs 28 °C: r = 0.075, P = 0.53): a lineage with a steep slope at one temperature was no more likely to have a steep slope at another. Together, the absence of any shared directional change in slope and the lack of genetic correlation in slope between temperatures indicate that temperature alters individual lineages’ nutritional scaling idiosyncratically, with no common direction. This supports the hypothesis that thermal and nutritional scaling are largely governed by independent mechanisms

### The pattern of individual-level scaling relationships does not change with temperature

Our previous work demonstrated that the pattern of individual-level scaling relationships in a population determine how the population-level relationship should respond to selection. Although we did not find any consistent change in individual-level nutritional scaling across temperatures here, it remains possible that the pattern of individual-level scaling relationships is temperature sensitive. This pattern is broadly defined by the relationship between the median point of intersection (MPI) among individual-level scaling relationships, and the bivariate mean of the traits: if the bivariate mean and the MPI intersect, the pattern is ‘see-saw’; if the bivariate mean is closer to the origin than the MPI, the pattern is ‘speedometer’; if the bivariate mean farther from the origin than the MPI, the pattern is ‘broomstick’ (**Figure 5A**). While the MPI and bivariate mean of the individual-level scaling relationships shifted with temperature, their position relative to each other did not change, suggesting that the pattern of individual-level scaling relationships is broomstick across all three temperatures (**Figure 5B**).

**Figure 5.**
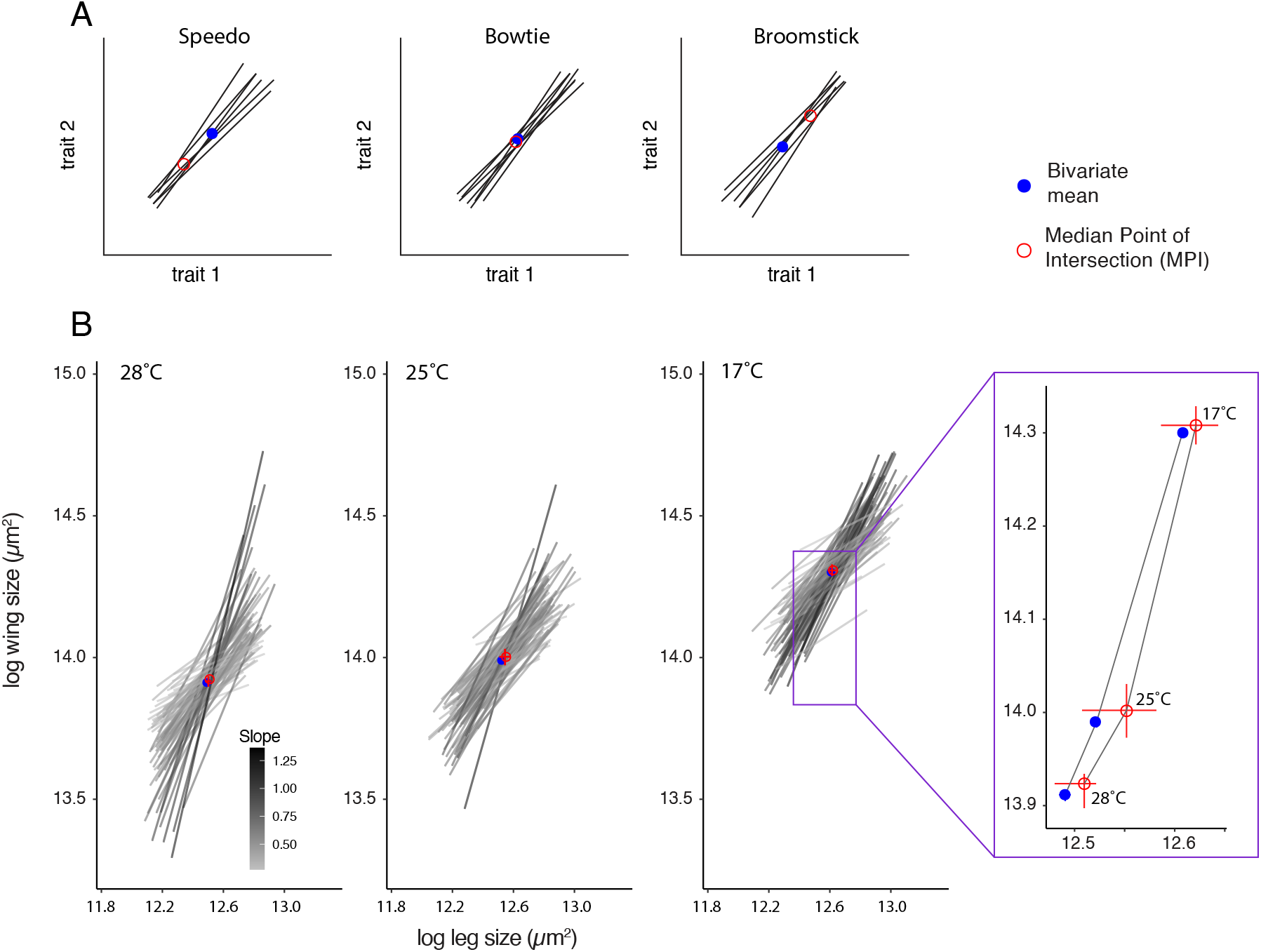
The pattern of the nutritional individual-level scaling relationships does not change with temperature. (A) The pattern of individual-level scaling relationships is categorized by the position of the median point of intersection (MPI, open red circle) among individual-level scaling relationships relative to the bivariate mean of trait size (closed blue circle), generating speedo, bowties or broomstick distributions. (B) At all three temperatures, the pattern of nutritional individual-level scaling relationships is more-or-less a bowtie. Inset show the position of the bivariate mean of wing and leg size and the median point of intersection at each temperature. Error bars are 95% HPD and are hidden in some cases.

### Discussion

The pattern of individual-level scaling relationships (ILSRs) in a population is typically hidden but is predicted to determine the response of the population-level scaling relationship (PLSR) to selection (Dreyer et al. 2016). Because different traits differ in their sensitivity to different environmental regulators of size, ILSRs may vary depending on the environmental factor generating size variation. What is unclear is whether this impacts the pattern of ILSRs in a population, and if so, the nature and consequence of this impact. Using temperature and nutrition as different generators of size variation, we found no genetic correlation between thermally- and nutritionally-induced size plasticity for a single trait, no genetic correlation between the slope of thermal and nutritional ILSRs, and no detectable change in the pattern of nutritional ILSRs under different thermal conditions. While there is some evidence that nutritional plasticity increases with temperature for both the wing and the leg, overall our data suggest that the developmental-genetic mechanisms that regulate scaling in response to temperature are largely independent of the mechanisms that regulate scaling in response to nutrition.

Historically, the study of phenotypic plasticity, both at proximate-developmental and ultimate-evolutionary levels, has been dominated primarily by focusing on the effect of a single environmental variable. Nevertheless, there is an increasing appreciation that organisms develop in multidimensional environments, generating multidimensional plastic responses (Relyea 2004; Westneat et al. 2019; Rodrigues and Beldade 2020; Hudak and Dybdahl 2023). The interaction of temperature with other environmental regulators of size, including nutrition, has perhaps received the most attention (Kingsolver et al. 2006; Stillwell et al. 2007; Lee and Roh 2010; Clissold and Simpson 2015; Kutz et al. 2019; Kim et al. 2020; Chakraborty et al. 2025). In *D. melanogaster*, the general trend appears to be an increase in nutritional plasticity of trait size with an increase in temperature (Chakraborty et al. 2023), with higher temperatures exacerbating the negative effects of a poor diet (Kutz et al. 2019). Our data are consistent with this observation, by indicating that nutritional plasticity increases with temperature in the both the wing and the leg in *D. melanogaster*.

While our data support an interaction between the effects of temperature and nutrition on trait size, we did not find evidence that this systematically influenced the slope of the wing-leg ILSR across temperatures. Further, the pattern of the nutritional ILSRs did not change between 17°C, 25°C and 28°C, with the MPI and the wing-leg bivariate mean maintaining their position relative to each other across temperatures. Consequently, although the position of nutritional ILSRs shifted toward the origin with increasing temperature, the pattern of these relationships was thermally insensitive (**Figure 5B**).

The constancy of the pattern of nutritional ILSRs across temperatures has important implications for the mode of selection that is predicted to alter the slope of PLSRs. Because the pattern of ILSRs for a population is predicted to determine the response of the PLSR to selection (Dreyer et al. 2016), if this pattern is thermally insensitive, the same form of selection will generate the same response regardless of the thermal environment at which individuals develop. For example, selection on increased or decreased relative wing size (wing:thorax ratio) in *D. melanogaster* increases and decreases the slope of the wing:thorax scaling relationship at 25°C, respectively (Robertson 1962). This is presumably because selection targets individuals that have correspondingly steep or shallow nutritional wing-thorax ILSRs (Dreyer et al. 2016). Our data suggest that the population will respond in the same way regardless of the temperature at which selection is being applied.

While the same form of selection on nutritional scaling may generate the same response at different temperatures, the selected genotypes responsible for this response may be different. This is because we did not detect a genetic correlation across temperature in the slope of a lineage’s ILSR – the lineage with the steepest wing:leg ILSR at 17°C did not have the steepest scaling relationship at 25°C. This was also evident as a significant random interaction between temperature and lineage on slopes, such that ILSR slopes changed with temperature, but differently for different lineages. The result is a higher-order (G×E_1_)×E_2_ interaction, in which the genetic architecture underlying variation in nutritional scaling (G×E_1_) – i.e. which genotypes contribute high-slope and low-slope phenotypes – is itself a function of the temperature (E_2_) at which nutritionally-induced size variation is generated. Consequently, the phenotypic response to selection at one temperature may have a different genetic basis for the same phenotypic response to selection produced at another temperature. Such (G×E_1_)×E_2_ interactions substantially complicate predictions of how traits respond to selection in heterogeneous environments (Sgrò and Hoffmann 2004; Stinchcombe et al. 2012; Westneat et al. 2019).

There are several caveats to the interpretation of our data. First, we studied only males. The slope and intercept of the wing:body and leg: body nutritional ILSRs are slightly but significantly different for females and males (Wilcox et al. 2023). It is possible, therefore, that the slope of wing:leg ILSR is genetically correlated across temperatures in females but not in males. The pattern of nutritional ILSRs is also slightly different in males than females (Wilcox et al. 2023), and this pattern may also change with temperature in females but not in males. However, the slope of the wing:body and leg:body ILSRs are genetically correlated between the sexes, so our findings in males may likely extend to females. Regardless, additional experiments are needed to explore the effect of sex on the pattern of ILSRs in *D. melanogaster*, and how this influences the effects of selection on ILSRs and the consequences for the PLSR.

A second caveat is that we did not explicitly impose variation in nutrition within our temperature treatments (17°C, 25°C and 28°C), in contrast to our experiments exploring nutritional scaling at 22°C. Although we found scaling within thermal treatments echoed the nutritional scaling relationship among genotypes, suggesting that size variation is due to access to nutrition, size variation was more limited within thermal treatments. This reduces the precision of the MA regressions to estimate the slopes of individual-level scaling relationships, expanding the confidence limits around our estimates. This in turn reduces the power of our analyses to capture genetic correlation between slopes, and may explain why we did not observe a genetic correlation in the slope of ILSRs across temperatures. Future experiments directly manipulating developmental nutrition at temperatures other than 22°C are necessary to more systematically test the hypothesis that ILSRs change with temperature.

Finally, our data were collected using isogenic lineages, reared under constant thermal conditions. Natural populations, however, comprise heterogenic individuals that experience fluctuating thermal conditions during development. Our original model of scaling evolution incorporated genetic heterogeneity among individual-levels, but did not include the (random) effects of temperature on scaling. Future iterations of this model should incorporate multidimensional plastic responses and their potential effects on the response of scaling to selection.

## Conclusions

In conclusion, our data are consistent with previous studies that suggest that nutritional and thermal scaling relationships are generated by distinct developmental and physiological processes. Further, our data suggest the hypothesis that, even if the phenotypic response to selection on scaling is the same across environments, the genetic basis of this response differ. A compelling method to test this hypothesis would be to impose artificial selection to change the slope of nutritional scaling relationships at two different temperatures. If our hypothesis were correct, the populations should show the same response in their PLSR, but this response should be due to selection on different alleles, so that the populations diverge in their genetic composition even as they converge in their phenotypes. Whether this is because different alleles target the same genes, signaling pathways or physiological mechanisms in the two populations is an open question. More generally, our findings suggest that the genetic basis of plasticity and scaling evolution in heterogeneous environments cannot be inferred from heritability or selection response measured in any single environment. Because scaling is the substrate of morphological evolution, and because the genetic architecture of scaling appears to vary across environments, the evolution of morphology in nature may be both richer and less predictable than single-environment studies suggest.

## Author Contributions

Conceptualization, AWS and WAF; Methodology, AWS, WAF and SMG; Data Collection: SMG and ASW; Formal Analysis, AWS.; Writing – Original Draft Preparation, AWS.; Writing – Review & Editing, AWS, WAF, SMG and ASW; Funding Acquisition, AWS and WAF. All authors have read and agreed to the published version of the manuscript.

## Data Availability Statement

All data and the R code used to analyze them are provided on Dryad.

## Acknowledgments

This work was made possible through the assistance of undergraduate members of the Shingleton and Frankino labs, who reared and measured the flies used in the study. Stocks obtained from the Bloomington Drosophila Stock Center (NIH P40OD018537) were used in this study.

## Supplementary Materials

### Supplementary Methods

The scaling relationship between wing and leg size across an environmental gradient can be estimated using an MA regression (Shingleton 2019). We calculated the slope of the scaling relationship using the mean wing and leg size for each lineage in two environmental conditions (fed – starved; 17°C – 25°C; 25°C – 28°C; 17°C – 28°C). The MA slope of the relationship is calculated as:

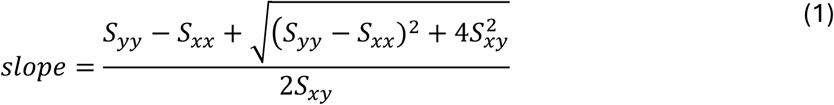

For two points (e.g. wing and leg size in fed versus starved flies):

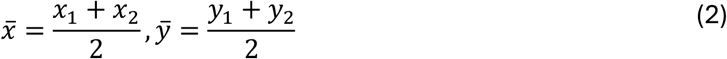

And

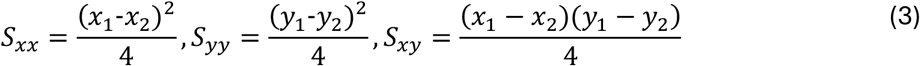

Inserting into Eqn 1:

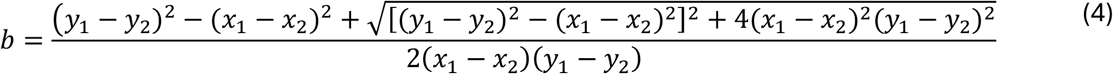

Simplifying:

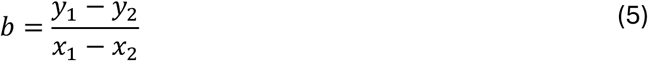

Thus the slope of the MA regression can be estimated as the ratio of the wing and leg plasticities acros

Shingleton AW. 2019. Symposium article: Which line to follow? The utility of different line-fitting methods to capture the mechanism of morphological scaling. Integr Comp Biol. 61:838 https://academic.oup.com/icb/advance-article-abstract/doi/10.1093/icb/icz059/5497803?redirectedFrom=fulltext. https://doi.org/10.1093/icb/icz059

## Supplementary Figures

**Supplementary Figure S1.**
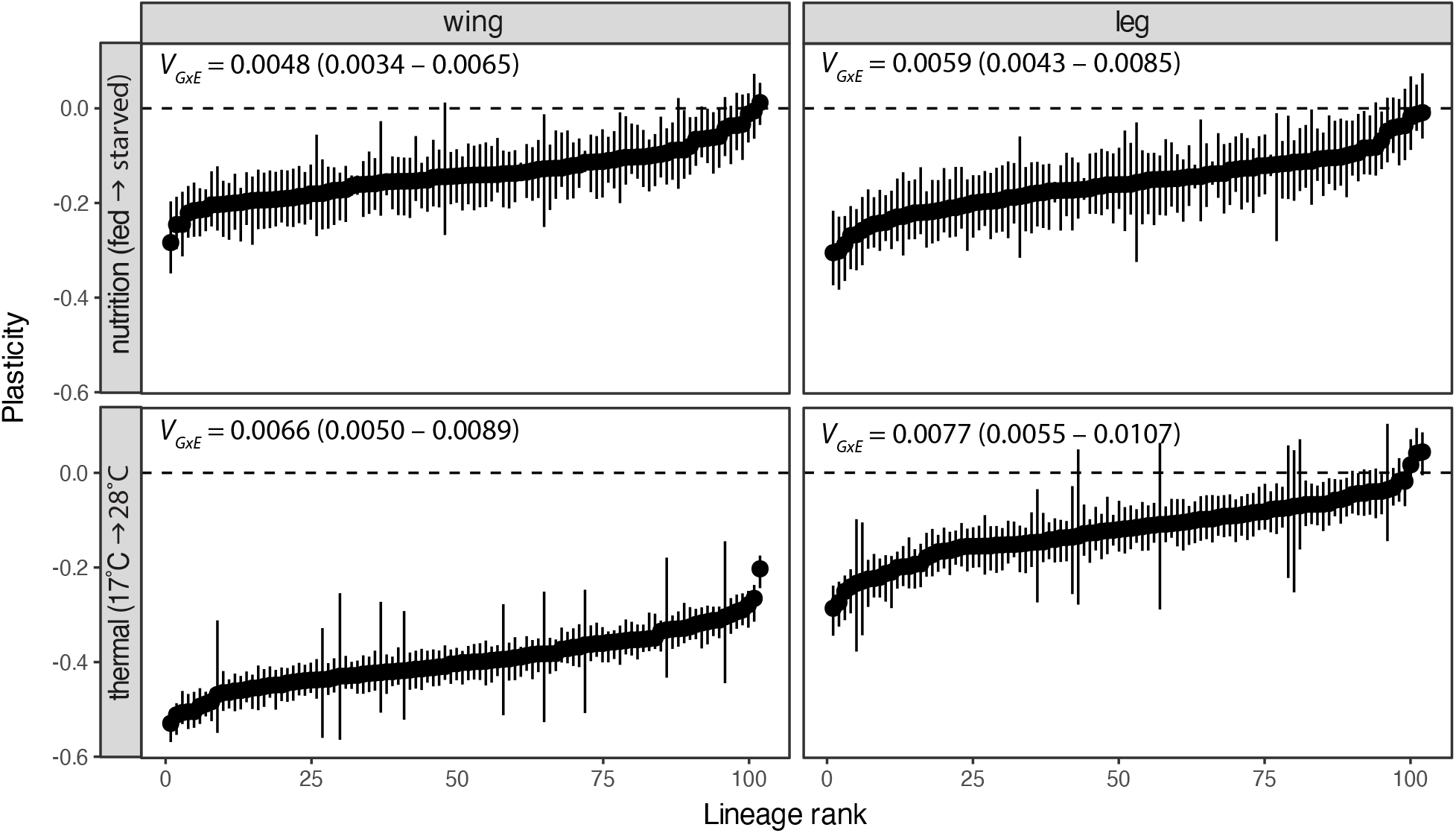
Plasticity in wing and leg size in response to nutrition (fed vs. starved) and temperature (17°C vs 28°C) among *Drosophila* lineages, ranked within each panel from most to least plastic. Points show the posterior mode with 95% HPD intervals. Almost all values are negative, reflecting reduced size under low nutrition or high temperature. *V*_*g*_ = modal genetic variance with 95% HPD intervals.

**Supplementary Figure S2.**
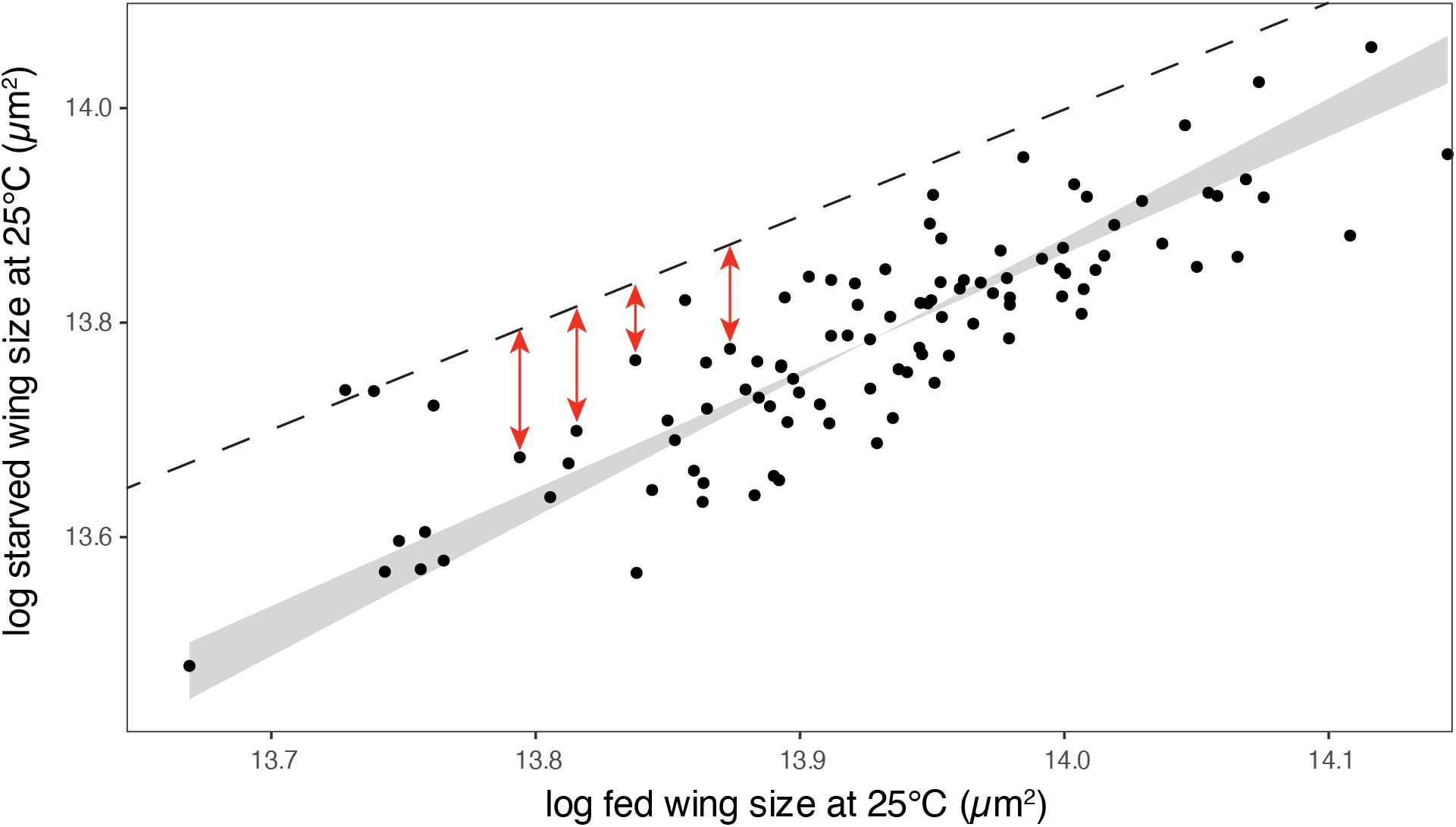
The genetic correlation among lineages of wing size in fed and starved flies. The modal genetic correlation is 0.8144 (95% HPD interval: 0.7675–0.8463) consistent with genetic variation in nutritional plasticity. Broken line represents where wing size is the same in both fed and starved flies and red arrows show lineage-specific plasticity. Because the slope of the regression is >1 (1.2034, 95% HPD interval:1.0948–1.2994), plasticity decreases as overall wing size increases. This would be true even if the genetic correlation were 1, indicating that a genetic correlation of < 1 is a suYicient but not necessary indicator of genetic variation in plasticity (GxE). Gray shading shows the 95% HPD interval for the MA regression of starved on fed wing size.

**Supplementary Figure S3.**
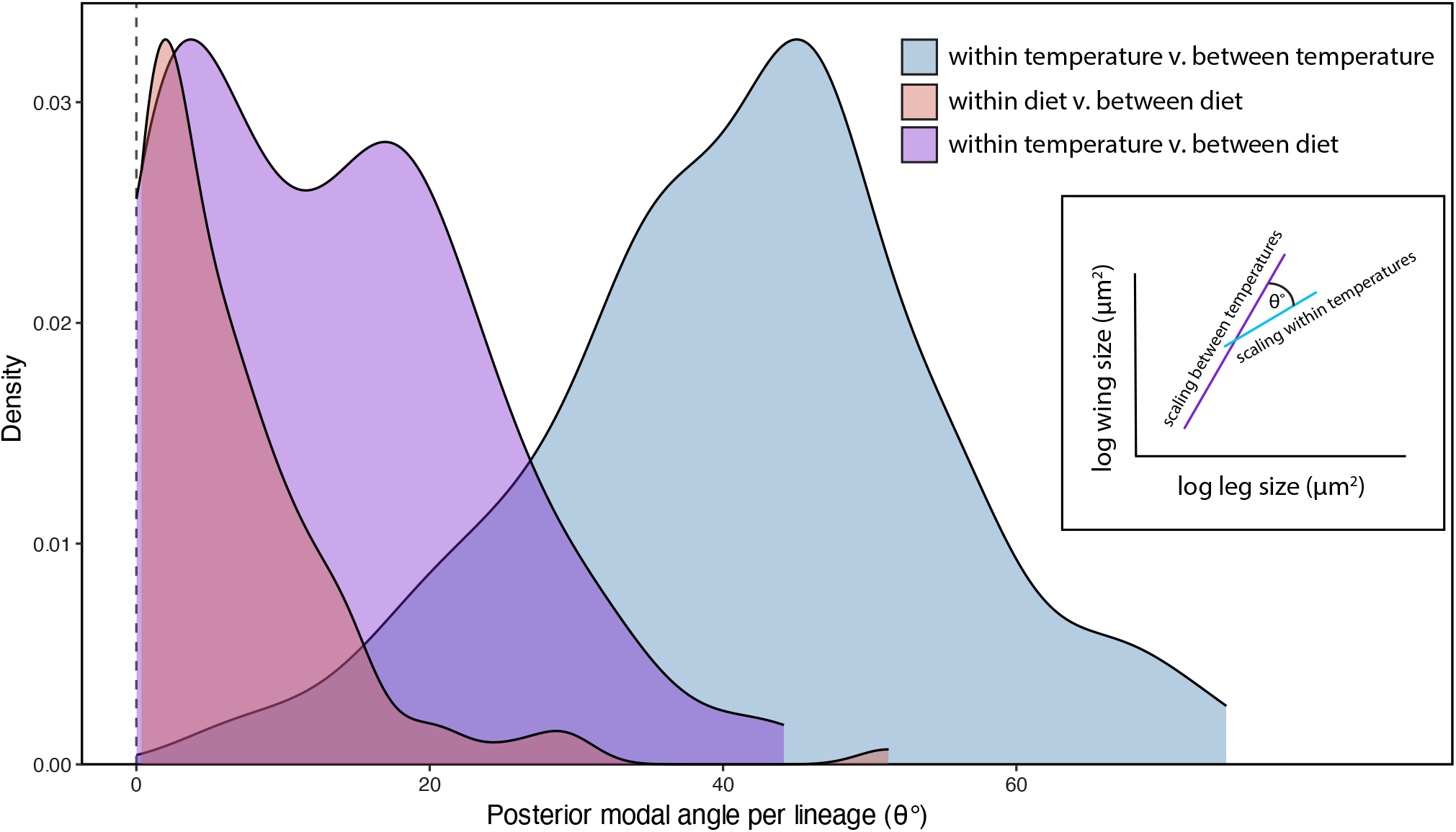
Within-temperature scaling relationships align with nutritional scaling relationships rather than thermal scaling relationships across lineages. For each lineage, we extracted two axes: Axis 1: the within-temperature scaling relationship, defined as the first eigenvector of the within-temperature wing-leg covariance, averaged across temperatures; Axis 2: the thermal/nutritional scaling relationship, defined as the first eigenvector of the covariance of predicted mean wing and leg size across temperatures/diets from the posterior draw. We then quantified the angle between these two axes per lineage for each posterior draws (inset), which reflects the degree to which each lineage’s thermal/nutritional scaling relationship is aligned with its within-temperature scaling relationship. An angle near 0° indicates that within-temperature scaling relationship for a lineage aligns with that lineage’s thermal/nutritional scaling relationship. The distribution is the modal angle across draws for each lineage.

## Supplementary Tables

**Supplementary Table 1.** Genetic variances (diagonal, blue), genetic covariances (upper triangle, purple), and slopes of the MA regression^†^ of trait size in one environment on trait size in another across lineages (lower triangle, orange), for wing and leg size; values shown as posterior mode (95% HPD interval).

|  |  | 17 ° C | 25 ° C | 28 ° C | Fed | Starved |
| --- | --- | --- | --- | --- | --- | --- |
| Wing | 17 ° C | 0.0042<br>(0.0037, 0.0046) | 0.5714<br>(0.4852, 0.6350) | 0.5041<br>(0.3891, 0.5812) |  |  |
|  | 25 ° C | 1.3518<br>(1.1819, 1.4994) | 0.0056<br>(0.0052, 0.0064) | 0.6675<br>(0.6020, 0.7158) |  |  |
|  | 28 ° C | 1.0419<br>(1.0948, 1.2994) | 0.6892<br>(0.5782, 0.7682) | 0.0042<br>(0.0037, 0.0049) |  |  |
|  | Fed |  |  |  | 0.0090<br>(0.0081, 0.0098) | 0.8144<br>(0.7675, 0.8463) |
|  | Starved |  |  |  | 1.2034<br>(1.0948, 1.2994) | 0.0018<br>(0.0106, 0.0130) |
| Leg | 17 ° C | 0.0054<br>(0.0050, 0.0062) | 0.6161<br>(0.5582, 0.6745) | 0.4318<br>(0.3319, 0.5251) |  |  |
|  | 25 ° C | 1.0913<br>(0.9692, 1.2462) | 0.0061<br>(0.0056, 0.0069) | 0.6586<br>(0.5987, 0.7248) |  |  |
|  | 28 ° C | 0.6454<br>(0.4884, 0.7759) | 0.6892<br>(0.5782, 0.7759) | 0.0036<br>(0.0032, 0.0043) |  |  |
|  | Fed |  |  |  | 0.0129<br>(0.0119, 0.0143) | 0.8125<br>(0.7683, 0.8476) |
|  | Starved |  |  |  | 1.0846<br>(0.9676, 1.1579) | 0.014<br>(0.0130, 0.0160) |

<sup>†</sup> Slope of the major axis regression of trait size in the row environment ( $y$ ) on trait size in the column environment ( $x$ ), across lineages. For each regression, the trait size in the column environment ( $x$ ) is typically larger than the trait size in the row environment ( $y$ ). Thus if the slope is $> 1$ , the plasticity increases with overall trait size, while if slope is $< 1$ , plasticity decreases with overall trait size (see Supplementary Figure 1).

**Supplementary Table 2.** Genetic correlation among plasticities within and between traits.

| | Comparison | $r^1$ | 95% HPD<sup>2</sup> |
| --- | --- | --- | --- |
| Wing vs Leg<br>(same condition) | Nutritional plasticity | **0.85** | **(0.79, 0.89)** |
|  | Thermal plasticity (17→25 °C) | **0.63** | **(0.53, 0.68)** |
|  | Thermal plasticity (25→28 °C) | **0.58** | **(0.49, 0.67)** |
|  | Thermal plasticity (17→28 °C) | **0.50** | **(0.35, 0.56)** |
| Within wing | Thermal (17→25 °C) vs (25→28 °C) | **-0.46** | **(-0.55, -0.35)** |
|  | Nutrition vs thermal (17→25 °C) | 0.12 | (-0.01, 0.22) |
|  | Nutrition vs thermal (25→28 °C) | -0.08 | (-0.19, 0.06) |
|  | Nutrition vs thermal (17→28 °C) | 0.06 | (-0.08, 0.19) |
| Within leg | Thermal (17→25 °C) vs (25→28 °C) | **-0.36** | **(-0.47, -0.21)** |
|  | Nutrition vs thermal (17→25 °C) | 0.00 | (-0.09, 0.16) |
|  | Nutrition vs thermal (25→28 °C) | -0.03 | (-0.14, 0.12) |
|  | Nutrition vs thermal (17→28 °C) | 0.00 | (-0.11, 0.15) |

<sup>1</sup> Correlation coefficient<sup>2</sup> 95% HPD. Intervals that exclude 0 are shown in **bold****Supplementary Table 3:** Genetic correlation among slopes of ILSRs generated by different environmental factors.

**Supplementary Table 3.**
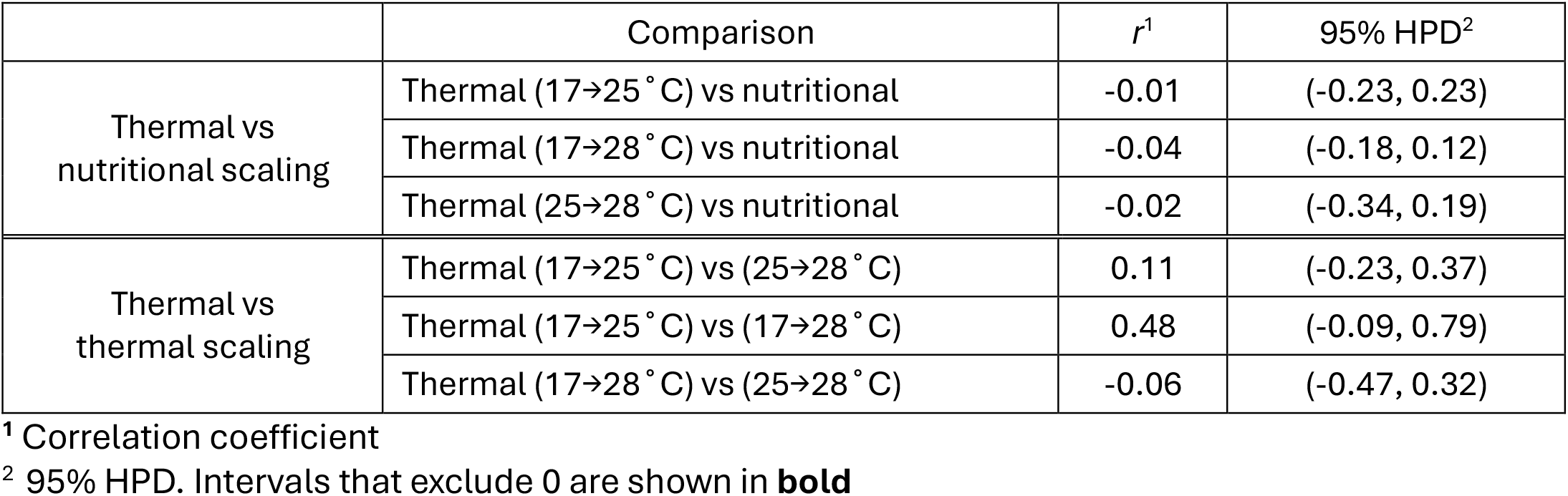
Genetic correlation among slopes of ILSRs generated by different environmental factors.

| | Comparison | $r^1$ | 95% HPD<sup>2</sup> |
| --- | --- | --- | --- |
| Thermal vs<br>nutritional scaling | Thermal (17→25 °C) vs nutritional | -0.01 | (-0.23, 0.23) |
|  | Thermal (17→28 °C) vs nutritional | -0.04 | (-0.18, 0.12) |
|  | Thermal (25→28 °C) vs nutritional | -0.02 | (-0.34, 0.19) |
| Thermal vs<br>thermal scaling | Thermal (17→25 °C) vs (25→28 °C) | 0.11 | (-0.23, 0.37) |
|  | Thermal (17→25 °C) vs (17→28 °C) | 0.48 | (-0.09, 0.79) |
|  | Thermal (17→28 °C) vs (25→28 °C) | -0.06 | (-0.47, 0.32) |

<sup>1</sup> Correlation coefficient<sup>2</sup> 95% HPD. Intervals that exclude 0 are shown in **bold**

